# IFNγ protects motor neurons from oxidative stress via enhanced global protein synthesis in FUS-associated Amyotrophic Lateral Sclerosis

**DOI:** 10.1101/2022.12.05.519150

**Authors:** Amanda Faria Assoni, Erika N. Guerrero, René Wardenaar, Danyllo Oliveira, Petra L. Bakker, Valdemir Melechco Carvalho, Oswaldo Keith Okamoto, Mayana Zatz, Floris Foijer

**Affiliations:** European Research Institute for the Biology of Ageing (ERIBA), University of Groningen, University Medical Center Groningen, Groningen, 9713 AV, the Netherlands; Instituto de Biociências, Universidade de São Paulo, São Paulo, SP 05508-900, Brazil; Department of Stem Cell Research, Gorgas Memorial Institute for Health Studies, Panama City, Republic of Panama; Division of Research and Development, Fleury Group, São Paulo, SP, 04344-070, Brazil

## Abstract

Amyotrophic lateral sclerosis type 6 (ALS6) is a familial subtype of ALS linked to Fused in Sarcoma (FUS) gene mutation. FUS mutations lead to decreased global protein synthesis, but the mechanism that drives this has not been established. Here, we used ALS6 patient-derived induced pluripotent stem cells (hIPSCs) to study the effect of the ALS6 FUS^R521H^ mutation on the translation machinery in motor neurons (MNs). We find, in agreement with findings of others, that protein synthesis is decreased in ALS6 MNs. Furthermore, ALS6 MNs are more sensitive to oxidative stress and display reduced expression of TGF-β and mTORC gene pathways when stressed. Finally, we show that IFN**γ** treatment reduces apoptosis of ALS6 MNs exposed to oxidative stress and partially restores the translation rates in ALS6 MNs. Overall, these findings suggest that a functional IFN**γ** response is important for FUS-mediated protein synthesis, possibly by FUS nuclear translocation in ALS6.

**Highlights and eTOC blurb:**

- ALS6 patient-derived motor neurons show decreased viability and reduced production of innate immune cytokines following oxidative stress
- FUS cytoplasmic localization coincides with decreased protein synthesis rates
- IFN**γ** treatment of ALS6 patient-derived motor neurons reduces apoptosis and ameliorates translation rates resulting from oxidative stress

## Introduction

Amyotrophic lateral sclerosis (ALS) is a late-onset neurodegenerative disease that affects motor neurons in the motor cortex, brainstem, and spinal cord (Charcot, 1869). It is characterized by muscle twitching, spasms, stiffness, weakness, and finally, muscle atrophy with a life expectancy of 3-5 years after diagnosis (Mathis et al., 2019). ALS is the most common motor neuron disease with an average prevalence and incidence of 4.42 and 1.59 per 1,000,000 population that increases with age (Xu et al., 2020). In ∼90% of the cases ALS is sporadic, but ∼10% of patients show familial mutations (Mathis et al., 2019. ALS subtypes are classified according to the affected gene, which include Superoxide Dismutase 1 (SOD1; ALS1), TAR DNA Binding Protein-43 (TDP-43; ALS10), Chromosome 9 Open Reading Frame (C9ORF72; ALS1), and Fused in Sarcoma (FUS; ALS6) (Guerrero et al., 2016), the latter leading to one of the most aggressive and early onset types of ALS (Vance et al., 2009).

FUS is a component of the heterogeneous nuclear ribonucleoprotein protein complex (hnRNP) and a DNA/RNA-binding protein involved in DNA damage repair, splicing, and multiple aspects of RNA metabolism16 (Wang et al., 2019, 2018). More than 50 different mutations in the FUS gene have so far been identified in ALS6 patients (Blair et al., 2010). While some mutations affect the N-terminal region, arginine-glycine-glycine box (RGG) and RNA recognition motif (RRM) region, most missense mutations affect the nuclear localization signal (NLS) domain in the C-terminus of the protein. These mutations result in cytoplasmic mislocalization and nuclear clearance of FUS, and yield an aggressive disease phenotype (Notaro et al., 2021; Vance et al., 2013). The most common FUS mutation affects arginine 521 (R to H, C, or G) (Shang and Huang, 2016). FUS mislocalization is not unique to ALS6, but also occurs in other familial forms of ALS and sporadic cases where FUS itself is not mutated (Tyzack et al., 2019). However, how FUS mutations lead to motor neuron death in ALS remains unclear.

Strategies to study the pathobiology of ALS are challenging since biopsies are associated with high cost and morbidity and do not provide a definitive pathological diagnosis (Nathani et al., 2021). Therefore, reliable models of the disease are key to elucidate the molecular mechanisms underlying ALS. In this study, we use iPSCs derived from ALS6 patients carrying a FUS^R521H^ mutation to generate motor neurons (MNs) to study ALS disease biology. As iPSCs rejuvenate as a result from reprogramming (Lapasset et al., 2011) and ALS symptoms present with ageing, we expose iPSC-derived motor neuros to oxidative stress to model ageing-associated effects (Albers and Flint Beal, 2000). Although patient-derived IPSCs present limitations in their ability to simulate age-associated traits, we consider this model crucial for understanding early phenotypes leading to developing therapeutics that may prevent disease progression. Our models reveal that MNs generated from ALS6 patient-derived iPSCs show aberrant cytoplasmic localization of FUS and decreased translation rates. Furthermore, ALS6 iPSC-derived MNs are more susceptible to oxidative stress-induced apoptosis than MNs differentiated from iPSCs generated from unaffected family members. This increased susceptibility coincides with decreased TGF-β and mTOR signaling and an altered cytokine landscape. Intriguingly, we find that supplementation of IFNγ to ALS MN-cultures reduces oxidative stress-induced apoptosis significantly, which coincides with improved of translation rates and nuclear FUS localization. Overall, our results show that IFNγ treatment reduces sensitivity to oxidative stress specifically of ALS6 MNs. While further work is required to understand how IFNγ restores FUS localization and impaired translation rates, our findings suggest that early-diagnosed ALS6 patients might benefit from IFNγ treatment to slow down disease progression.

## Results

### iPSC-derived MNs from ALS6 patients are susceptible to oxidative stress-induced apoptosis

To investigate how mutant FUS affects the biology of MNs, we generated IPSC lines from 2 ALS6 patients carrying the FUS^R521H^ alongside with 2 IPSC lines from phenotypically healthy relatives not carrying the mutation (from now on referred to as ‘control’). IPSCs were differentiated into mature MNs according to an established protocol (Du et al., 2015) (**Figure 1A**) followed by immunofluorescence (IF) stainings for HB9, MAP2, and motor neuron-specific markers Tuj1, Islet1 to confirm complete differentiation (**Figure 1B**). We then compared cell viability and apoptosis rates between control and ALS6 IPSCs, Neural Progenitor cells (NPS) and mature MNs, but found no significant differences (**Figure 1C, D**). As oxidative stress has been proposed as a driving factor in ALS (Richard and Maragakis, 2015), cells were treated with sodium arsenite (SA), a well-known inducer of oxidative stress through generation of reactive oxygen species (ROS) (Baron et al., 2013). ALS6 mature MNs were significantly more sensitive to SA treatment compared to healthy controls, evidenced by a lower viability (**Figure 1E, F**) and increased apoptosis following treatment (**Figure 1G, H**). We conclude that ALS6- and control iPSCs, NPCs and MNs have similar viability under unperturbed conditions, but that ALS6-derived MNs are more sensitive to oxidative stress induced by SA.

**Figure 1.**
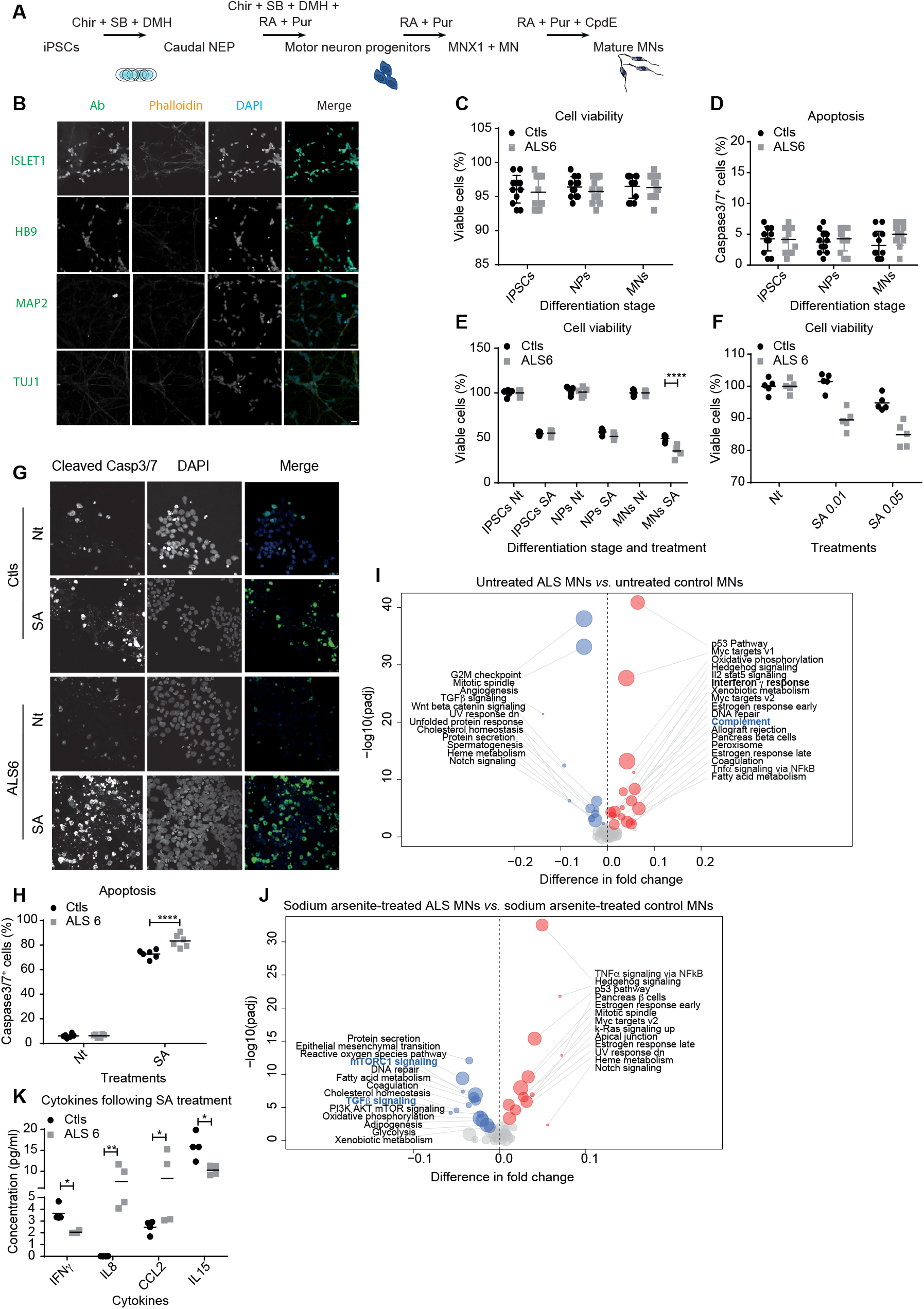
ALS motor neurons are more sensitive to oxidative stress than control motor neurons (MNs). **A**. Schematic representation of the differentiation protocol used to obtain MNs. **B**. Representative images from immunofluorescence stainings to characterize MNs following the differentiation protocol. Scale bar represents 20 μm. **C**. MTS assays of untreated IPSCs, NPs and MNs 24 hours following plating. **D**. Percentage of Caspase 3/7-positive untreated IPSCs, NPs and MNs 24h after plating **E, F**. MTS assays of control- (Nt) or SA-treated IPSCs, NPs and MNs (E) or control or ALS6 MNs treated with different doses of SA (F) 24h after plating. **G**. Representative images from immunofluorescence stainings for Caspase3/7 of control- (Nt) or SA-treated MNs. MNs were treated for 24 h. Scale bar represents 25 μm. **H**. Percentage of Caspase 3/7-positive control- (Nt) or SA-treated MNs. Cells were treated for 24 h. **I**. Volcano plot showing differentially regulated Hallmark pathways between untreated ALS and control MNs. **J**. Volcano plot showing differentially regulated Hallmark pathways between SA-treated ALS and control MNs **K**. Quantification of cytokines secreted by SA-treated MNs. Cells were treated for 24 h. * = p< 0.05; ** = p< 0.01; and *** = p< 0.001, two-way ANOVA with Tukey multiple comparison test; n = 8 per group.

### ALS6 motor neurons show altered expression of genes in the mTOR pathway and genes related to the innate immune response system

To explore why ALS6 MNs are more sensitive to oxidative stress than controls, we compared the transcriptomes of ALS6 and control MNs, either or not treated with SA, by RNA sequencing. Interestingly, ALS6 MNs showed increased innate immune system (*i*.*e*. “complement”) transcripts compared to healthy control MNs (**Figure 1I**), but only when not treated with SA. In contrast, SA-treated ALS6 MNs showed decreased expression of genes involved in TGF-β and mTORC signaling compared to SA-treated healthy controls (**Figure 1J**). As TGF-β is critical for the activation of cytokines (Cottrez and Groux, 2001; Hayashi et al., 2004), next we measured cytokines secreted by ALS6 MNs when exposed to oxidative stress using a cytokine multiplex assay. Indeed, we found that ALS6 MNs secrete more IL-8 and CCL2, and less INFγ and IL-15 when treated with SA (**Figure 1K**). In line with this, we also found that control MNs upregulate INFγ signaling following SA treatment (**Supplemental Figure 1A**), but ALS MNs fail to do so (**Supplemental Figure 1B**). These observations suggest that ALS6 MNs show an impaired response to oxidative stress involving decreased TGF-β and mTORC signaling and an altered cytokine landscape including reduced IFNγ.

### Global translation rates are decreased in ALS6 NPs and MNs

As ALS6 MNs displayed reduced expression of genes involved in mTORC signaling following SA treatment and mTORC and FUS both have known roles in protein synthesis (López-Erauskin et al., 2018; Ma and Blenis, 2009; Martin and Blenis, 2002), we next assessed protein translation in IPSCs, NPs or MNs from ALS6 patients and controls. To quantify translation rates, we used a puromycin (a tRNA mimic) incorporation assay, followed by quantification of incorporated puromycin by Western blot (WB) and IF using an anti-puromycin antibody, a commonly used assay to measure protein synthesis rates (**Figure 2A-C**) (Aviner, 2020). While protein synthesis rates were similar between all IPSC samples, untreated AL6 NPs and MNs displayed significantly decreased translation rates compared to untreated control NPs and MNs. This was not the case when comparing treated ALS6 to control MNs, possibly because translation rates were already low in ALS6 MNs to begin with (**Figure 2D, E**). We also tested whether transcription rates were affected in ALS6 cells, but found no overt differences compared to wild type cells in iPSCs, NPCs nor MNs (**Supplemental Figure 2A, B**). These observations indicate that untreated ALS MNs display a translation defect, in agreement with findings of others (de la Fuente and Emc, 2020; Kamelgarn et al., 2018; López-Erauskin et al., 2018).

**Figure 2.**
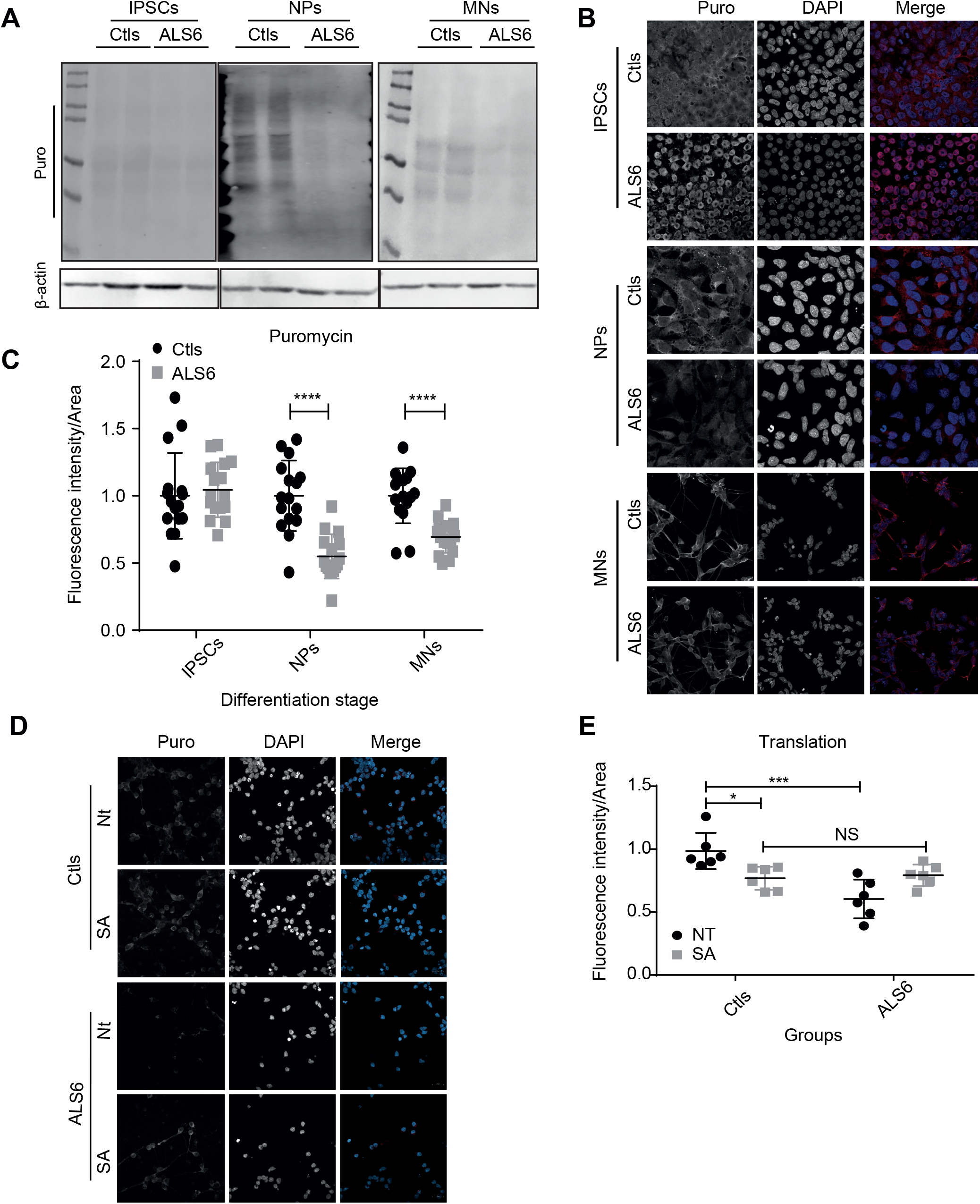
Global translation rates are decreased in ALS6 cells. **A**. Western blot with anti-puromycin antibody on total cell extract of IPSCs, NPs and MNs. **B**. Representative images of immunofluorescence stainings of incorporated puromycin into IPSCs, NPs and MNs. Scale bar for IPSCs represents 25 μm and 10 μm for NPs and MNs. **C**. Quantification of the relative intensity of puromycin incorporation in immunofluorescence stainings on IPSCs, NPs and MNs. **** = p<0.0001; two-way ANOVA with Sidak’s multiple comparison test; n = 16 per group. **D**. Representative images of immunofluorescence stainings of incorporated puromycin antibody into control- or SA-treated MNs treated. Scale bar represents 10 μm. **E**. Quantification of the relative intensity of puromycin incorporation from immunofluorescence stainingson control- or SA-treated MNs. * = p<0.05; *** = P<0.001, two-way ANOVA with Sidak’s multiple comparison test; n = 16 per group.

To determine whether the abnormalities in the ALS MNs are the result of a gain-of-function mutation in the FUS gene or the result of reduced levels of functional FUS protein, we next modulated wild type FUS or mutant FUS^R521H^ levels in control and ALS6 MNs by transient overexpression and RNA interference (**Supplemental Figure 2C, D**). We found that overexpression of wild type FUS in ALS cells nor downregulation of FUS in control MNs affected protein translation rates (**Supplemental Figure 2E-H**). However, when we transiently overexpressed mutant FUS^R521H^ in control MNs, we did observe significantly decreased translation rates (**Supplemental Figure 2 I, J**). These data strongly suggest that the decreased translation rates observed in ALS MNs result from a gain-of-function mutation and not reduced levels of FUS functional protein. Overall, these observations suggest that ALS6 MNs are more susceptible to apoptosis due to decreased protein synthesis rates caused by a R521H gain-of-function mutation in the FUS gene.

### FUS is mislocalized into the cytoplasm in ALS6 MNs and interacts with proteins of the translational machinery in the cytoplasm

To better understand how mutant FUS leads to decreased translation, we next compared (mutant) FUS interactors between control and ALS6 MNs. For this, we co-immunoprecipitated (co-IPed) endogenous wild type and/or mutant FUS from ALS6 and control MNs followed by proteomic shotgun identification of the protein interactors (**Figure 3A-C**). Interestingly, co-IPed (mutant) FUS in ALS6 samples had a higher number of protein interactors compared to FUS isolates from control samples (**Figure 3A; Supplemental Table 1**). Interestingly, FUS binding partners unique to the ALS6 isolates were enriched for proteins involved in translation initiation that are localized in the cytoplasm (**Figure 3B, C**). Note that the increased number of interactors in ALS6 MNs was not the result of increased expression of FUS as we observed no significant differences in FUS protein levels between wild type and ALS6 cells across differentiation stages (**Figure 3D, E**).

**Figure 3.**
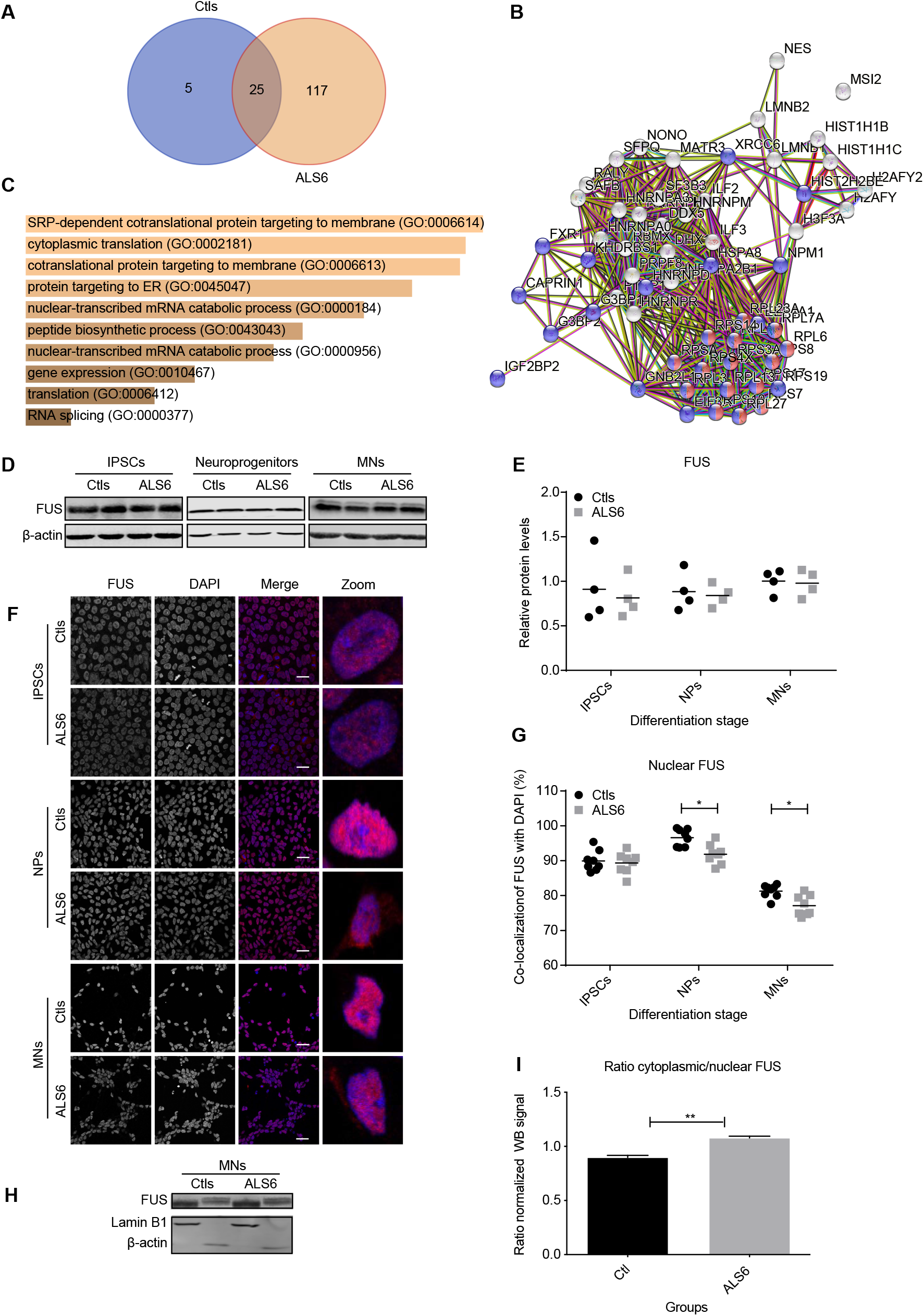
FUS localizes into the cytoplasm in ALS6 cells and interacts with cytoplasmic proteins of the translational machinery. **A**. Venn diagram of proteins interacting with FUS in each sample type. **B**. String association diagram of FUS interacting proteins unique to ALS6 samples. **C**. Panther bar graph of enriched pathways in FUS interacting proteins unique to ALS6 samples. **D**. Western blot of total FUS protein levels in IPSC, NP, and MNs samples. **E**. Densitometric quantification of the total amount of FUS in each sample relative to β-actin (n=4 each group). **F**. Representative images of immunofluorescence stainings of FUS showing nuclear localization in control and nuclear localization in all cells and partial and cytoplasmic localization in ALS MNs Scale bar represents 10 μm. **G**. Quantification of FUS localization from immunofluorescence stainings as sown in (f) in control and ALS6 IPSCs, NPs and MNs. * = p<0.05; two-way ANOVA with Sidak’s multiple comparison test; n = 8 per group). **H**. Western blot for FUS protein in nuclear and cytoplasmic fractions of control and ALS6 MNs. Lamin serves as a nuclear control and Actin as a cytoplasmic control. **I**. Quantification of the ration between cytoplasmic and nuclear FUS determined from Western blots (n = 2 samples for each condition) as shown in (H).

Even though FUS is known to shuttle between nucleus and cytoplasm in healthy cells, in ALS, FUS becomes predominantly cytoplasmic in MNs, ultimately leading to FUS aggregates (Deshpande et al., 2019). Indeed, when we assessed FUS localization in ALS6 iPSCs, NPs and MNs, we found that FUS levels were reduced in the nuclei and modestly increased in the cytoplasm of NPCs and MNs, but not iPSCs (**Figure 3F, G**). Wild type iPSCs and NPCs mostly showed nuclear FUS as expected. Control MNs showed a reduction in nuclear FUS compared to iPSCs and NPCs, which was further reduced in ALS6 MNs, evidenced from IF stainings as well as Western blots of cytoplasmic and nuclear fractions of MNs (**Figure 3F-I**).

As FUS has also been reported to be mislocalized in the cytoplasm of postmortem neurons from sporadic ALS patients (Tyzack et al., 2019), we next assessed FUS localization in MNs generated from iPSCs from patients with other ALS subtypes, including MNs from ALSp1, ALSp2, ALSp3 and ALSp4 patients. Also here, we detected reduced nuclear FUS levels compared to healthy controls (**Supplemental Figure 3A, B**), suggesting that aberrant FUS localization is not unique to ALS6 patients. We also compared translation rates of these cells to control cells and found that translation rates were significantly decreased in MNs-derived from ALSp1-4 compared to healthy controls (**Supplemental Figure 3C, D**). Together these findings suggest that aberrant cytoplasmic localization of FUS is common among ALS subtypes and leads to promiscuous binding of FUS to cytoplasmic proteins with a role in translation.

### IFNγ treatment rescues motor neuron viability following sodium arsenite treatment

As cytokine measurements showed reduced secreted IFNγ in ALS6 MNs treated with SA, and we recently found that inflammatory cytokines can protect cells from stress-induced cell death (Hong et al., 2022), we tested whether this was also true for ALS6 MNs. For this, we supplemented SA-treated ALS6 MNs with IFNγ and found that this indeed increased viability of ALS6 MNs to levels similar to SA-treated control MNs (**Figure 4A, B**). Furthermore, IFNγ treatment reduced the fraction of apoptotic ALS6 MNs following SA exposure (**Figure 4C, D**). Importantly, we also found that IFNγ treatment resulted in an increased IFNγ transcriptional response in SA-treated ALS MNs (**Figure 4E**), indicating that IFNγ treatment was sufficient to rescue the impaired IFNγ response observed in SA-treated ALS MNs (compare **Supplemental Figure 1B** to **Figure 4E**).

**Figure 4.**
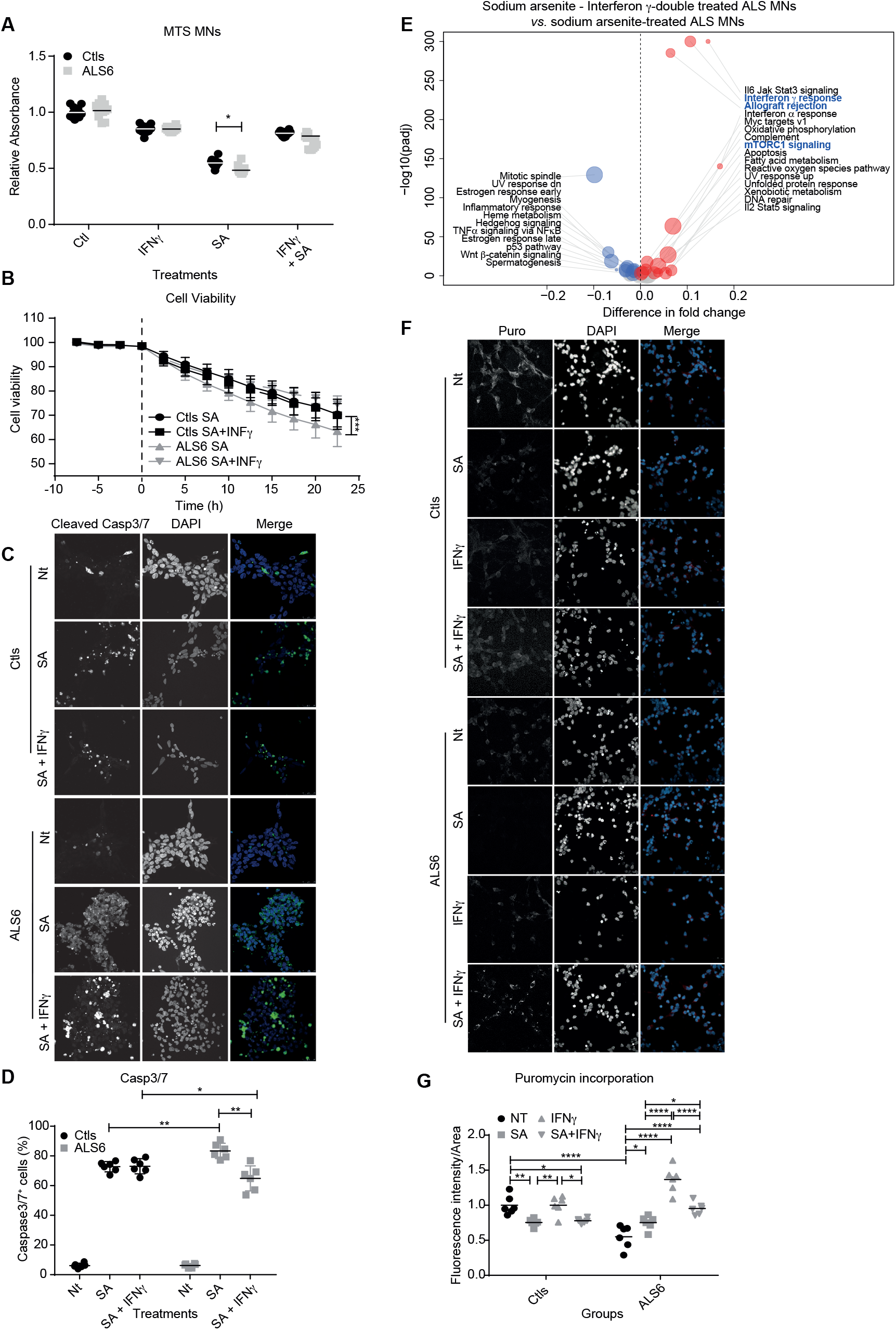
INFγ treatment rescues oxidative stress-induced decreased viability of ALS6 MNs. **A**. Cell viability of control (Nt), SA, INFγ or combination-treated control and ALS6 MNs assessed by the MTS assays. *** = p<0.001, two-way ANOVA with Tukey multiple comparison test; n = 8 per group. **B**. Cell viability of control (Nt), SA, INFγ or combination-treated control and ALS6 MNs assessed by xCELLigence real-time cell analysis system. *** = p<0.001, two-way ANOVA with Tukey multiple comparison test; n = 8 per group. **C**. Representative images of immunofluorescence stainings for Caspase3/7 on control (Nt), SA, INFγ or combination-treated MNs. Cells were treated for 24 h. Scale bar represents 25 μm. **D**. Percentage of Caspase 3/7-positive control (Nt), SA, INFγ or combination-treated MNs. Cells were treated for 24 h. * = p<0.05; ** = p<0.01; and *** = p<0.001, two-way ANOVA with Tukey multiple comparison test; n = 8 per group. **E**. Volcano plot showing differentially regulated Hallmark pathways between SA + INFγ ALS and SA-treated MNs. **F**. Representative images of immunofluorescence staining for incorporated puromycin in MNs for the four following treatments: Control (Nt), IFNγ, SA, and SA + IFNγ. Scale bar represents 20 μm. **G**. Relative intensity of puromycin incorporation from immunofluorescence stainings for puromycin incorporation into MNs for the four following treatments: Control (Nt), IFNγ, SA, and SA + IFNγ. * = p<0.05; ** = p<0.01; and *** = p<0.001, two-way ANOVA with Tukey multiple comparison test; n = 8 per group.

As ALS6 MNs showed impaired translation rates and these coincide with increased sensitivity to SA treatment, we next determined whether IFNγ treatment improved translation rates in ALS6 MNs. Translation rates were lower in ALS6 cells not exposed to SA, in agreement with our own findings (**Figure 2**) and those of others (Kamelgarn et al., 2018; López-Erauskin et al., 2018). Intriguingly, while IFNγ treatment did not affect translation rates in treated and untreated control MNs, it significantly increased translation rates in untreated as well as SA-treated ALS6 MNs (**Figure 4F, G**), in line with our observation that IFNγ treatment lead to increased expression of genes involved in mTORC signaling (**Figure 4E**). Finally, we investigated the effect of INFγ on FUS localization and found that INFγ treatment increased nuclear localization of FUS almost to levels in control MNs (**Supplemental Figure 4A, B**). Altogether, these results indicate that IFNγ treatment of ALS6 MNs improves their disease phenotype and decreases their sensitivity to SA, suggesting that ALS6 patients could benefit from IFNγ treatment to delay disease onset and/or progression.

## Discussion

In this study, we identify IFNγ as a potential compound to protect ALS6 motor neurons from oxidative stress. FUS is ubiquitously expressed and shuttles between the cytoplasm and nucleus in different cell types, but in healthy neurons FUS is mainly localized in the nucleus (Ishigaki and Sobue, 2018). In addition to the role of FUS in transcription and splicing, FUS also plays a role as an mRNA transporter between nucleus and cytoplasm, for instance in neuronal dendrites to transport mRNA into dendritic spines, which is critical for neuronal maturation (Fujii, 2005). Cytoplasmic mislocalization and thus reduced nuclear FUS is a well-known hallmark of ALS. Indeed, more than 50% of FUS mutations are clustered near the C-terminal NLS domain, underscoring the importance of defective nuclear import and the resulting enhanced cytoplasmic localization phenotype for the pathobiology of ALS (Guerrero et al., 2016).

To investigate the effect of the FUS^R521H^ mutation, common to ALS6, we used patient-derived iPSC models and generated motor neurons to study the disease biology in a highly physiological setting. Interestingly, although we find modest transcriptional differences between healthy control and ALS6 MNs, the viability of untreated ALS6 MNs is identical to control MNs, suggesting that both cultures manage to achieve homeostasis. However, as reprogramming somatic cells into iPSCs coincides with cellular rejuvenation, our MN model could well be recapitulating young MNs, while ALS(6) symptoms typically present later in life. To model age-associated damage, we therefore added an oxidative stress component to our model system by exposing the cells to sodium arsenite (SA) (Akbari et al., 2022; Baron et al., 2013). Furthermore, increased oxidative stress and ROS accumulation has been extensively reported as a hallmark of ALS as ROS are byproducts of cellular respiration as neurons have higher metabolic rates and transcriptional activity (Vandoorne et al., 2018). Indeed, we find that ALS6 MNs carrying the FUSR^521H^ mutation are more sensitive to SA treatment than matched control MNs consistent with previous studies (Wang et al., 2018).

We used RNA-seq to comprehensively evaluate the impact of FUS^R521H^ mutation on ROS sensitivity in untreated and SA-treated MNs. We find increased expression of innate immune system components in untreated mutant cells, suggesting that ALS6 MNs exhibit intrinsic inflammation, in accordance with several lines of evidence suggesting toxic effects of the inflammatory response during disease progression (de Boer et al., 2014; Haidet-Phillips et al., 2011; Heitzer et al., 2016; Mirzaei et al., 2022; Potenza et al., 2016). We also find that ALS6 MNs display a lower expression of genes in the TGF-β signaling pathway following SA-treatment. This is quite interesting as TGF-β signaling has been described as neuroprotective following oxidative stress (Dhandapani and Brann, 2003; Galbiati et al., 2020; Prehn et al., 1993) and suggests that the failure of ALS to induce TGF-β following SA might be related to the increased sensitivity to SA-induced oxidative stress. Furthermore, cytokines measurements show that SA-treated ALS6 MNs secrete more IL-8 and CCL2, and less IL-15 and IFNγ compared to SA-treated control MNs. This is interesting, as other studies have reported elevated levels of IL-15 and IFNγ in serum and CSF of sporadic ALS patients (Liu et al., 2015; Rentzos et al., 2010). It is important to note here that we quantified cytokine secretion from MNs directly, while these other studies measured cytokines in serum, which might not fully represent what happens nearby the motor neurons. However, future work using iPSC-derived MNs modeling other subtypes of ALS should reveal whether the decreased secretion of IL-16 and IFNγ by MNs is specific to FUS^R521H^ patients or a feature more general to (sporadic) ALS MNs.

It is still not fully clear whether the FUS^R521H^ mutation and its cytoplasmic localization results from a gain-of-toxicity or loss-of-function effect in the FUS protein. Our results are consistent with previous studies suggesting that the FUSR^521H^ mutation leads to a gain of function that impairs protein translation (Jun et al., 2017; Kamelgarn et al., 2018; Sephtona et al., 2014; Udagawa et al., 2015) as we have identified a large number of FUS protein-protein interactors unique to the FUS^R521H^ protein. We find that these interactors have a function in translation initiation and are localized in the cytoplasm. Furthermore, studies in humanized mice expressing FUS^R521H^ showed an activated stress response and impaired local intra-axonal protein synthesis in hippocampal neurons and sciatic nerves without nuclear loss of function (López-Erauskin et al., 2018)].

We also observed that a small fraction of the FUS^R521H^ protein localizes to the cytoplasm, consistent with studies showing mild cytoplasmic localization of mutant FUS^R521G^, FUS^R521H^ and FUS^R521C^ protein (Vance et al., 2013), suggesting that mild cytoplasmic localization of FUS is sufficient to disrupt cellular protein synthesis homeostasis. We show that this cytoplasmic localization in MNs is not unique to ALS MNs with mutations in FUS but also occurs in MNs derived from IPSCs of patients with familial forms of ALS linked to mutations in the VAPB and VRK1 genes. In agreement with this, these MNs also displayed decreased translational rates. Therefore, our findings are consistent with other studies that suggest that mutant FUS represses translation through association with polyribosomes (Sévigny et al., 2020).

When we treated ALS6 MNs with IFNγ following the induction of ROS, we found that IFNγ - treated FUS^R521H^ MNs are less sensitive to SA-induced ROS. Intriguingly, IFNγ treatment also led to decreased cytoplasmic localization of FUS and increased protein translation rates, suggesting that IFNγ treatment decreases the impact of ROS somewhere upstream in the stress signaling cascade. There is conflicting evidence regarding the relationship between neurons and IFNs, as both protective as well as detrimental effects of IFNs have been suggested for ALS neurons (Beghi et al., 2000; Hashioka et al., 2015; Lubina-Dąbrowska et al., 2017; Poutiainen et al., 1994). For instance, it has recently been shown that IFNγ stimulation of adult human astrocytes yields neurotoxicity *in vitro* (Hashioka et al., 2015). Furthermore, studies with high doses of INFα in ALS patients result in cognitive decline, memory and psychomotor impairment, neurotoxicity and other side effects (Färkkilä et al., 1984; Iivanainen et al., 1985; Poutiainen et al., 1994). Conversely, both IFN-β1b and IFN-β1a treatment were found to inhibit the pro-inflammatory cytokines (IL-6, IL-1β, TNF-α and IFN-γ), increase the myelin protein level in the brain cortex, and improve the neurological status of experimental autoimmune encephalomyelitis rats (Lubina-Dąbrowska et al., 2017). Finally, INFβ treatment in ALS patients has not shown any significant difference between patients given INFβ-1a and patients given placebo (Beghi et al., 2000). Therefore, while our work suggests a positive effect of IFNγ treatment, at least for ALS6 patients, further work is needed to compare the exact role of individual interferon signaling components between various ALS and control neuronal cell types.

Altogether, our work indicates that ALS patient-derived MNs are more sensitive to SA-induced oxidative stress compared to control MNs and that IFNγ improves survival, specifically of FUS^R521H^ MNs. This IFNγ-mediated rescue coincides with increased translation rates and decreased cytoplasmic localization of mutant FUS. While further work is required to disentangle the exact underlying molecular mechanism of this rescue, our findings might suggest that IFNγ, when dosed timely and appropriately, could protect motor neurons in ALS6 patients and might thus delay disease onset or progression. However, before testing this in patients, further work is required, such as validating the neuro-protective effects of IFNγ treatment in ALS6-mouse models (López-Erauskin et al., 2018) or, possibly in patient-derived MN models that better recapitulate the effects of natural aging, for instance by direct conversion of neurons from fibroblasts isolated from patients presenting ALS symptoms (Tang et al., 2017), or by overexpressing aging-inducing factors such as Progerin (Miller et al., 2013).

## Experimental procedures

### Patient material

Fibroblasts from patients and unaffected family members were collected at the Centro de Pesquisas sobre o Genoma Humano e Células-Tronco. Sampling was approved by the Comitê de Ética em Pesquisa do Instituto de Biociências da Universidade de São Paulo – IBUSP #5464/ Certificate CAAE # 20108413.4.0000.5464.

### Cellular reprogramming and motor neuron differentiation

Fibroblasts were reprogrammed using CytoTune™-iPS 2.0 Sendai Reprogramming Kit (Thermo) as per manufacturer instructions. iPSCs were cultured in Essential 8 medium (Thermo). Motor neuron (MN’s) differentiation was performed as previously described (Du et al., 2015). Briefly, iPSCs were cultured in NB medium containing DMEM/F12, Neurobasal medium, N2, B27 and Glutamax (Thermo). Differentiation was performed using a two-step protocol: neural induction and caudalization and ventralization to obtain motor neuron progenitors (MNP’s). For this, iPSCs were cultured for six days in NB medium with Dorsomorphin (2 μM) (Sigma), SB431542 (2 μM) (Sigma), CHIR99021 (3 μM) (Tocris) and Ascorbic acid (0.1 mM) (Sigma) for neural induction, followed by a six-day culture in NB medium containing Dorsomorphin (2 μM) (Sigma), SB431542 (2 μM) (Sigma), CHIR99021 (1 μM) (Tocris), retinoic acid (0.1 μM) (Sigma), Ascorbic acid (0.1 mM) (Sigma) and Purmorphamine (0.5 μM) (Tocris) for caudalization and ventrilization of MNP’s. MNP’s were seeded in 60 mm^2^ plates coated with Matrigel (Corning) for motor neuron differentiation with NB medium containing retinoic acid (0.5 μM) (Sigma), Purmorphamine (0.1 μM) (Tocris) and Ascorbic acid (0.1 mM) (Sigma) for six days. For neural maturation Compound E (0.1 μM) (Calbiochem) was added to NB medium with retinoic acid (0.5 μM) (Sigma), Purmorphamine (0.1 μM) (Tocris) and Ascorbic acid (0.1 mM) (Sigma). All cells were cultured at 37 °C and 5% CO2 in a humidified incubator and routinely checked for mycoplasma infection.

### Immunofluorescence

Motor neurons were fixed with 3.7% formaldehyde (Sigma) in 1x PBS (Gibco) for 20 minutes at room temperature followed by a 30-minute permeabilization step using 0.1% Triton X-100 (Thermo) in 5% bovine serum albumin (BSA) (Sigma). Protein labeling was done by incubation with appropriate primary antibodies as indicated (Supplementary Table S1) in 5% BSA (Sigma) at 4°C overnight, followed by incubation with the appropriate fluorescent secondary antibody for 45 minutes. Slides were washed 2 times with 1x PBS (Gibco) and counterstained with 1 μg/mL DAPI (Thermo) for 2 minutes to label nuclei and mounted with VectaShield (Vector Laboratories). Images were taken using a confocal microscope (Zeiss LSM 800). Quantification was performed using Cell Profiler 3.0 as previously described (Danielson et al., 2021; McQuin et al., 2018).

### Immunoblotting

Motor neurons were harvested by accutase (Gibco) dissociation and lysed with elution buffer (150 mM NaCl - Sigma, 0.1% NP-40 - Sigma, 5 mM EDTA-Sigma, 50 mM HEPES - Sigma pH7.5 and protease inhibitor cocktail-Sigma) for 15 minutes at 4°C and centrifuged for 10 minutes at 300xg at 4°C to obtain cell lysates. Proteins were separated on a 10% polyacrylamide gel and transferred onto a polyvinylidene difluoride (PVDF) membrane (Sigma). Membranes were blocked in Odyssey blocking buffer (Li-cor Biosciences) for 60 minutes at 4°C. Membranes were incubated with primary antibodies overnight at 4°C as indicated. Following primary antibody incubation membranes were washed 3 times with 0.1% Tween 20 (Sigma) in 1x PBS (Gibco) followed by incubation with the appropriate secondary antibody for 1 hour at room temperature. Western blots were detected using the Odyssey imaging system (Li-cor Biosciences). Quantification of the bands was performed using Image studio lite software (Li-cor Biosciences).

### Cytokine measurements

Cytokines were quantified in media isolated from MN cultures using the Bio-Plex Pro Human Cytokine kit (Biorad, USA) according to the manufacturer’s instructions. MNs were plated at a density of 5×10^5^ cells/well and treated with SA (5μM) (Sigma), IFNγ (50ng/mL) (Peprotech), or a combination for 24h prior to measurements.

### Co-Immunoprecipitation

Co-IPs for FUS were performed using a Immunoprecipitation Kit Dynabeads Protein A/G (ThermoFisher) according to manufacturer’s protocol. Briefly, protein A/G beads were incubated with 10μL of the relevant antibody or mouse IgG as a control. Cells were washed once with PBS and lysed in 500 μL/well lysis buffer containing 20 mM Tris pH 8.0, 10% glycerol, 135 mM NaCl, 0.5% NP-40, and protease inhibitors (Complete, EDTA-free, Roche) for 15 min on ice. After harvesting, cells were centrifuged at 16,100 x g for 5 minutes at 4°C to remove cell debris. The supernatant was incubated with antibody-conjugated beads for 1 hour at 4°C. Following 3 washing steps with wash buffer, beads were taken up in RapiGest (Waters) for 10 minutes at 65°C.

### Liquid Chromatography coupled to tandem mass spectrometry

Samples were digested using RapiGest (Waters) as a surfactant agent. Proteins were reduced with DTT (Sigma) and alkylated with iodoacetamide (Sigma). Samples were further digested with trypsin proteomic level (Promega) in enzyme : protein ratio of 1:50. Samples were processed by nanoACQUITY system with a binary pump, an auxiliary pump and a sampler. Desalted and concentrated peptides were captured in a Symmetry C18 column (2nm x 180mm, 5 um) in a mobile phase (composed of water with 0.1% trifluoroacetic acid) at 15uL/ min flow rate for 5 minutes. Further, peptides were separated in a analytical HSSC18 column (75 μm × 150 mm, 1.7 μm) by elution with a linear gradient of 2% DMSO (Sigma) in water with 0.1% formic acid (Sigma) and 5% DMSO in acetonitrile (LiChrosolv) with 0.1% formic acid. The proportion of the organic solution was increased from 0% to 60% in 80 minutes.

The chromatographic system was directly coupled to a hybrid quadrupole orbitrap tandem mass spectrometer Q-Exactive (Thermo Scientific), equipped with a Nano Flex source. Acquisition of spectral data was obtained by a data dependent top-15 method in which the spectrometer chooses dynamically the most abundant not-yet sequenced precursor ions from a survey scan from 390 to 1650 m/z (except for the monocharged and those with charges exceeding 7) at 70,000 (at m/z 200) of resolution and AGC target 5 e 6. Sequencing was achieved dissociating the precursor ion with normalized collision energy of 35, resolution equal to 17,500 and AGC target of 5 e 4.

Acquired data were processed using MaxQuant 1.4.0.8 proteomics data analysis workflow 27. Protein identification was performed by the Andromeda search tool using the database of the human proteome UniProtKB (SWISSPROT November 2017). The following criteria were applied for protein identification: 1) a maximum of two incomplete cleavages by trypsin, 2) fixed modification by carbamidomethylation of cysteines, and 3) variable modification by acetylation of the N-terminal portion and methionine oxidation. Quantification was based on the LFQ 28 label-free method.

### Puromycin incorporation assay

Cells were incubated with or without 20μM puromycin (Invitrogen) and fixed for immunofluorescence or lysed for immunoblotting.

### Subcellular Fractionation

Cytoplasmic/nuclear fraction separation was performed by dissolving 3×10^6^ cells in 200 μL of cold hypotonic buffer (10 mM HEPES−NaOH pH 7.9, 10 mM KCl, 1.5 mM MgCl2, 1 mM DTT, and protease and phosphatase inhibitors – all compounds from Sigma). Cells were incubated on ice and inverted every 5 minutes for 30 minutes, followed by centrifugation for 10 minutes at 400 x g at 4°C. The supernatant was taken as the cytoplasmic fraction and the pellet was washed 3 times with cold hypotonic buffer. Pellets were dissolved in cold 100 mM Tris-HCl (pH 9.0) containing 12 mM SDC (Sigma), 12 mM SLS (Sigma), and protease and phosphatase inhibitors, and collected as the nuclear fraction. Both fractions were subjected to SDS-page and western blot.

### EU Labeling

Cells were incubated with 10 μM EU (5-ethynyl-uridine; Click-It EU Alexa Fluor 488 Imaging Kit; Life Technologies) for 30 min. Cell nuclei were labeled with DAPI (Sigma) at a concentration of 5 μg/ml for 5 min. Images were taken using a confocal microscope (Zeiss LSM 800). Quantification was performed using Cell Profiler 3.0 as previously described (Danielson et al., 2021; McQuin et al., 2018).

### MTS assay

MTS assays were performed according to manufacturer’s instructions. Briefly, 15,000 motor neurons were plated per well and treated with IFNγ (50ng/mL), SA (5μM) and for the time indicated at relevant figures. MTS reagent (10 μl) (Promega) was added directly to the wells and incubated at 37°C for 4 hours. Absorbance was measured at 490 nm on a SpectraMax Plus 384 reader (Molecular Devices; Sunnyvale, Ca).

Cell viability assay kill curves were quantified by the impedance-based xCELLigence real-time cell analysis system (ACEA Biosciences, San Diego, CA, USA). For this, 50 μL of cell culture media was added to each 96 wells of the E-Plate 96 PET (ACEA Biosciences) for background reading. Subsequently, 50 μL of cell suspension containing 15,000 cells was added to each well and placed inside the xCELLigence incubator. Cells were treated with IFNγ (50ng/mL) (Peprotech), SA (5μM) (Sigma) and impedance reflecting changes in cell adhesion and cell death were measured every 15 min for 24 hours. Data are presented as changes of impedance (‘Cell Index’) over time, according to the manufacturer’s instruction.

### RNA Sequencing

RNA was isolated using a RNeasy mini kit (Qiagen). RNA quality and quantity of the total RNA was assessed by the 2100 Bioanalyzer using a Nano chip (Agilent, Santa Clara, CA). Total RNA samples having a RIN>8 were subjected to library generation. Strand-specific libraries were generated using the TruSeq Stranded mRNA sample preparation kit (Illumina Inc., San Diego, RS-122-2101/2), according to the manufacturer’s instructions (Illumina, Part #15031047 Rev. E).

Polyadenylated RNA from intact total RNA was purified using oligo-dT beads. Following purification, the RNA was fragmented, random primed and reverse transcribed using SuperScript II Reverse Transcriptase (Invitrogen, part 72 #18064-014) with the addition of Actinomycin D. Second strand synthesis was performed using Polymerase I and RNaseH with replacement of dTTP for dUTP. The generated cDNA fragments were 3′-end adenylated and ligated to Illumina Paired-end sequencing adapters and subsequently amplified by 12 cycles of polymerase chain reaction. Sequencing libraries were analyzed on a 2100 Bioanalyzer using a 7500 chip (Agilent, Santa Clara, CA), diluted and pooled equimolar into a 10 nM sequencing stock solution. Illumina TruSeq mRNA libraries were sequenced at a resolution of 50 base single reads on a HiSeq2000 using V3 chemistry (Illumina Inc., San Diego). Resulting reads were trimmed using Cutadapt (version 1.12) to remove any remaining adapter sequences, filtering reads shorter than 20 bp after trimming to ensure efficient mapping. The trimmed reads were aligned to the GRCm38 reference genome using STAR (version 2.5.2b). QC statistics from Fastqc (version 0.11.5) and the above-mentioned tools were collected and summarized using Multiqc (version 0.8). Gene expression counts were generated by featureCounts (version 1.5.0-post3), using gene definitions from Ensembl GRCm38 version 76. Normalized expression values were obtained by correcting for differences in sequencing depth between samples using DESeq median-of-ratios approach, and subsequent log-transformation of the normalized counts.

### Statistical analyses

Experiments were performed in at least biological triplicates as stated in the figures. Data were analyzed by one-way and two-way ANOVA followed by Bonferroni post hoc testing. A two-tailed unpaired t-test was used for pairwise comparison. GraphPad Prism software was used to perform all statistical analysis (version 6.0 GraphPad Software Inc.). Quantification of data is presented as mean ± standard error of the mean (SEM), and p-value thresholds are presented as: * = p<0.05; ** = p<0.01; *** = p<0.001; and **** = p<0.0001.

### Data availability

All RNA sequencing data has been deposited at ArrayExpress under accession number E-MTAB-12422. All proteomic data has been deposited at PRIDE under accession number PXD038042.

## Author contributions

**Amanda F. Assoni:** Conceptualization; data curation; formal analysis; validation; investigation; visualization; methodology; writing – original draft; writing – review and editing. **Erika N Guerrero**: data curation; formal analysis; visualization; writing – original draft; writing – review and editing. **René Wardenaar**: Data curation; formal analysis; investigation; methodology. **Danyllo Oliveira:** Investigation, visualization; methodology. **Petra L. Bakker**: Investigation, methodology, validation. **Valdemir Melechco Carvalho:** Investigation, methodology. **Oswaldo Keith Okamoto:** Resources; data curation; formal analysis; supervision; funding acquisition. **Mayana Zatz:** Conceptualization; supervision; resources. **Floris Foijer:** Conceptualization; resources; data curation; formal analysis; supervision; funding acquisition; investigation; visualization; methodology; writing – original draft; project administration; writing – review and editing.

## Acknowledgments

We thank the people in the Foijer and Zatz labs for fruitful discussions. This work was supported by the Fundação de Amparo à Pesquisa do Estado de São Paulo (FAPESP), Conselho Nacional de Desenvolvimento Científico e Tecnológico (CNPq) and an Abel Tasman fellowship to AA awarded by the University of Groningen.

## Figure Legends

**Supplemental Figure 1.**
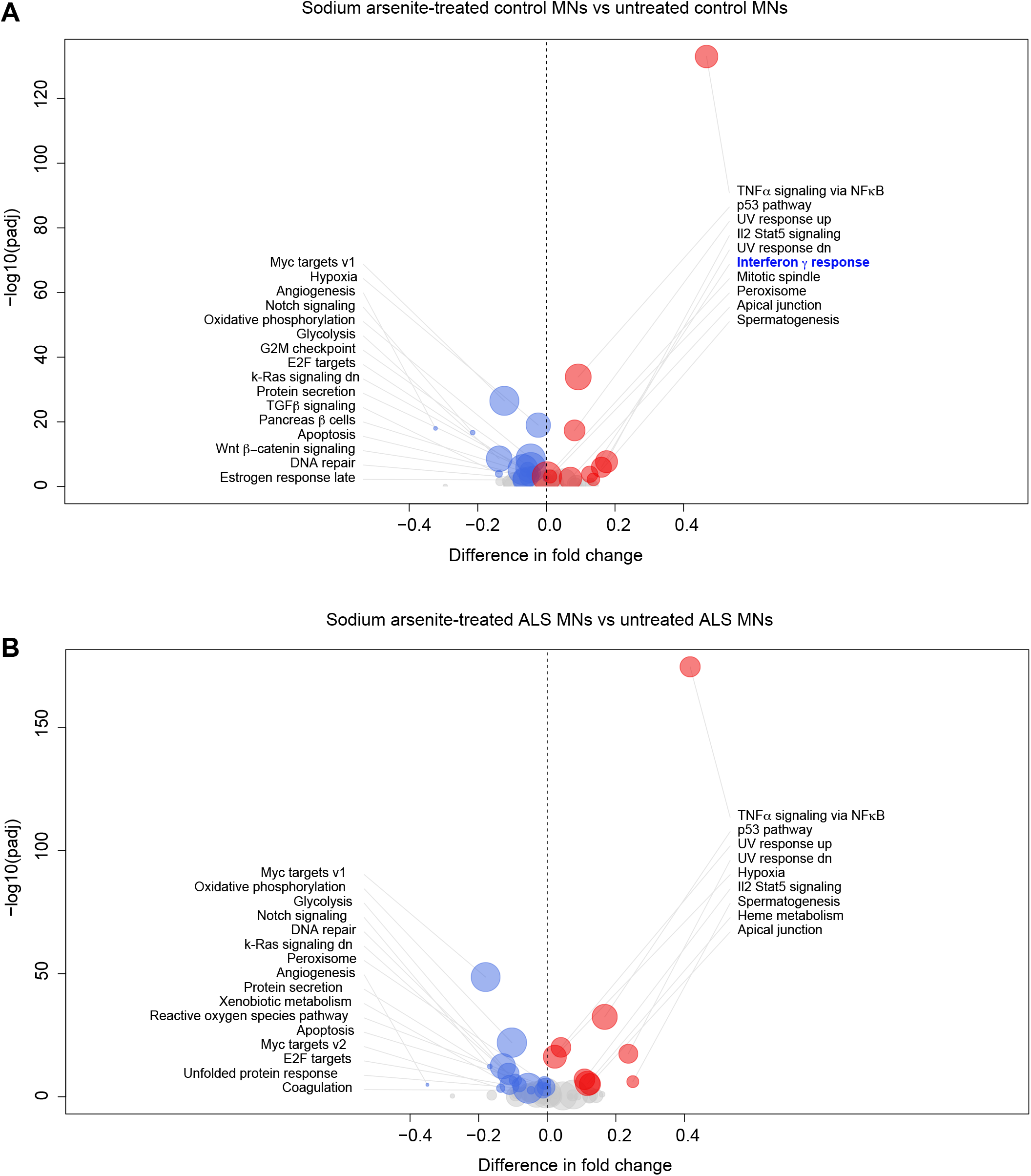
Related to Figure 1. Sodium arsenite induces an IFNγ response in control MNs, but not ALS6 MNs. **A**. Volcano plot showing differentially regulated Hallmark pathways between mock- and SA-treated control MNs. **B**. Volcano plot showing differentially regulated Hallmark pathways between mock- and SA-treated ALS6 MNs.

**Supplemental Figure 2.**
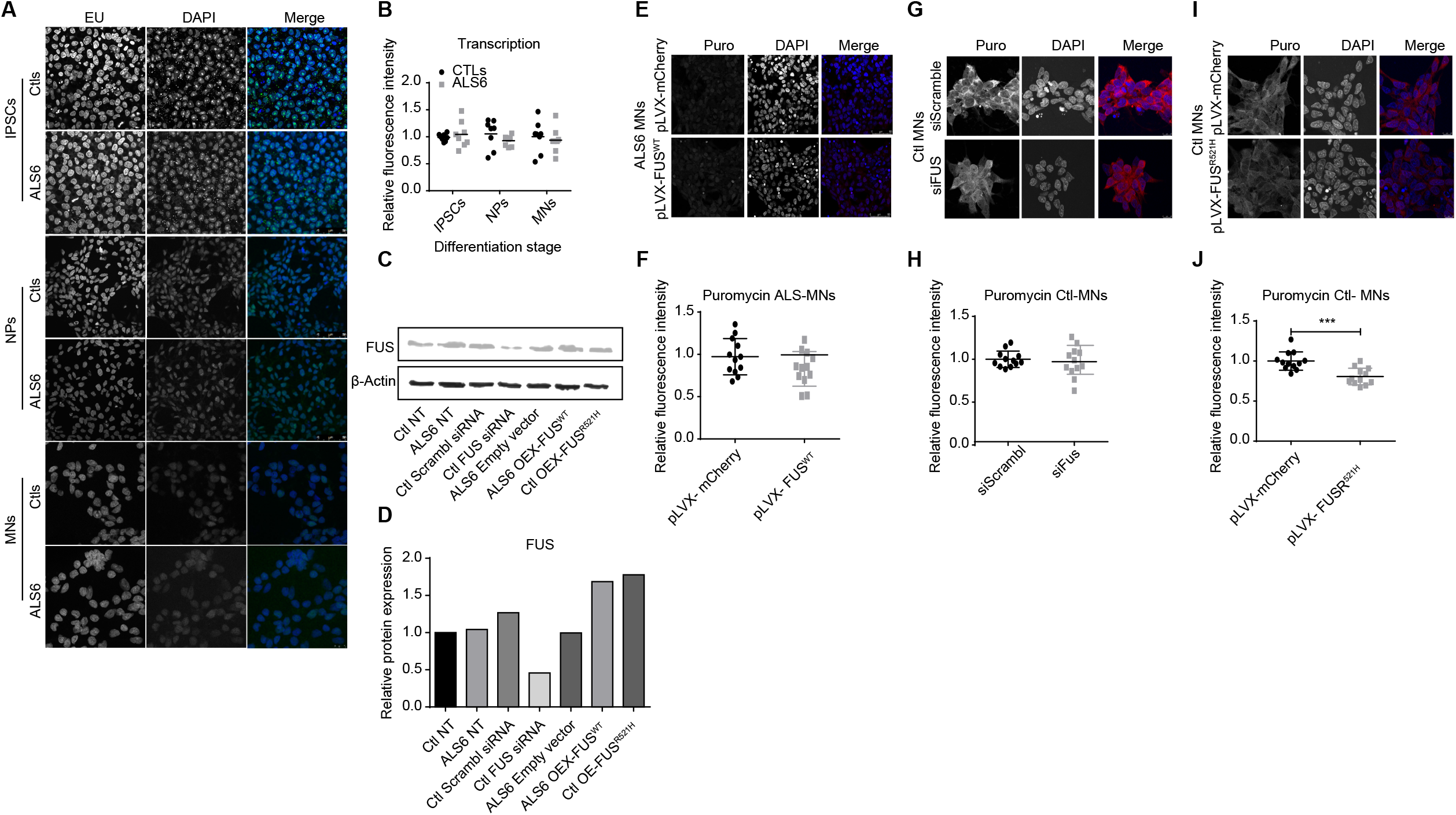
Related to Figures 1 and 2. Presence of mutant FUS changes global translation rates in control MNs, which cannot be explained by altered transcription. **A**. Differentially expressed biological processes between untreated ALS6 and control MNs. **B**. Differentially expressed biological processes between SA-treated ALS6 and control MNs. **C**. Western blots of total FUS protein in MNs transfected with FUS siRNA or a vector for FUS overexpression (OE). **D**. Densitometric quantification of the total amount of FUS protein in each sample relative to β-actin (n=1 each group). **E**. Representative images of immunofluorescence stainings for incorporated puromycin in ALS6 MNs transiently transfected with the overexpression plasmids pLVX-mCherry or pLVX-mCherry-wild type FUS. Scale bar represents 10 μm. **F**. Relative fluorescence intensity in images of immuno-fluorescence for puromycin incorporation in ALS6 MNs transiently transfected with the pLVX-mCherry or pLVX-mCherry-wild type FUS overexpression plasmids (n=12). **G**. Representative images of immunofluorescence stainings for puromycin incorporation into control MNs transiently transfected with siRNA targeting endogenous FUS, or a scrambled control. Scale bar represents 10 μm. **H**. Relative fluorescence intensity in images of immuno-fluorescence for puromycin incorporation in control MNs transiently transfected with siRNAs targeting endogenous FUS, or a scrambled control (n=12). **I**. Representative images of immunofluorescence stainings for incorporated puromycin into control MNs transiently transfected with pLVX-mCherry or pLVX-mCherry-FUS^R521H^ overexpression plasmids. Scale bar represents 10 μm. **J**. Relative fluorescence intensity in images of immuno-fluorescence for puromycin incorporation in control MNs transiently transfected with the overexpression plasmids pLVX-mCherry or pLVX-mCherry-FUS^R521H^. *** = p<0.001; Mann Whitney test; n=12 per group. **K**. Representative images immune-fluorescence stainings for 5-Ethynyl-uridine (5-EU) incorporation to measure RNA transcription in IPSCs, NPs and MNs. Scale bars represent 50μm (iPSCs), NPs (50μm) and MNs (10 μm). **L**. Relative intensity of EU incorporation from immunofluorescence stainings on IPSCs, NPs and MNs (n=8).

**Supplemental Figure 3.**
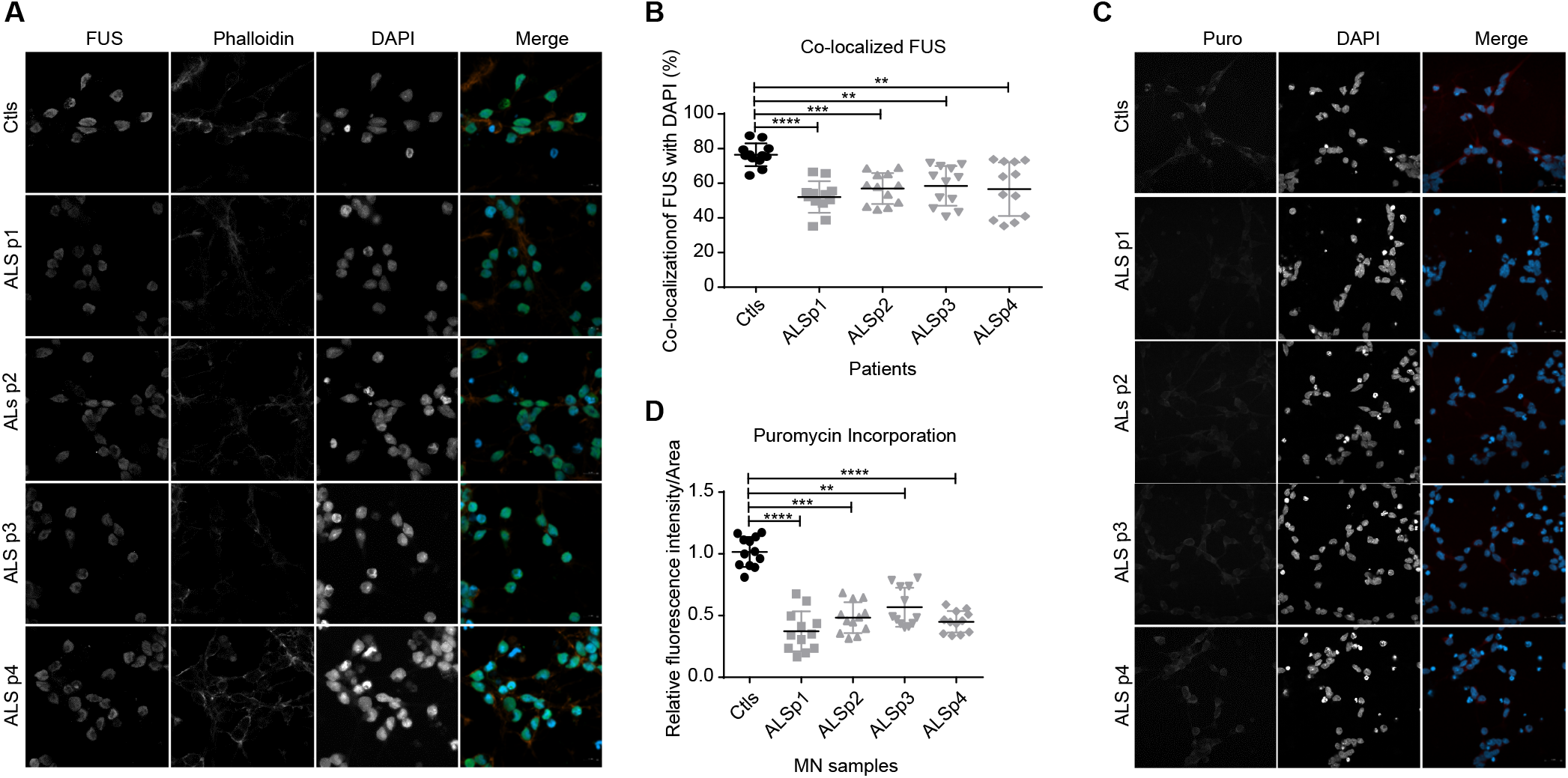
Related to Figure 3. MNs from other ALS subtypes show similar characteristics to ALS6 MNs. **A**. Representative images of immunofluorescence stainings for FUS protein to determine localization in IPSCs-derived MNs from control and ALS patients. Scale bar represents 10 μm. **B**. Quantification of FUS localization from FUS immunofluorescence stainings on Ctls and ALS patient IPSCs-derived MNs. ** = p<0.01; *** = p<0.001; and **** = p<0.0001, Kruskal-Wallis test and Dunn’s multiple comparisons, n=12 each. **C**. Representative images of immunofluorescence stainings for incorporated puromycin into IPSCs-derived MNs from control and ALS patients. Scale bar represents 20 μm. **D**. Relative level of puromycin incorporation from immunofluorescence stainings on control and ALS patient IPSCs-derived MNs. ** = p<0.01; *** = p<0.001; and **** = p<0.0001, Kruskal-Wallis test and Dunn’s multiple comparisons, n= 12 each).

**Supplemental Figure 4.**
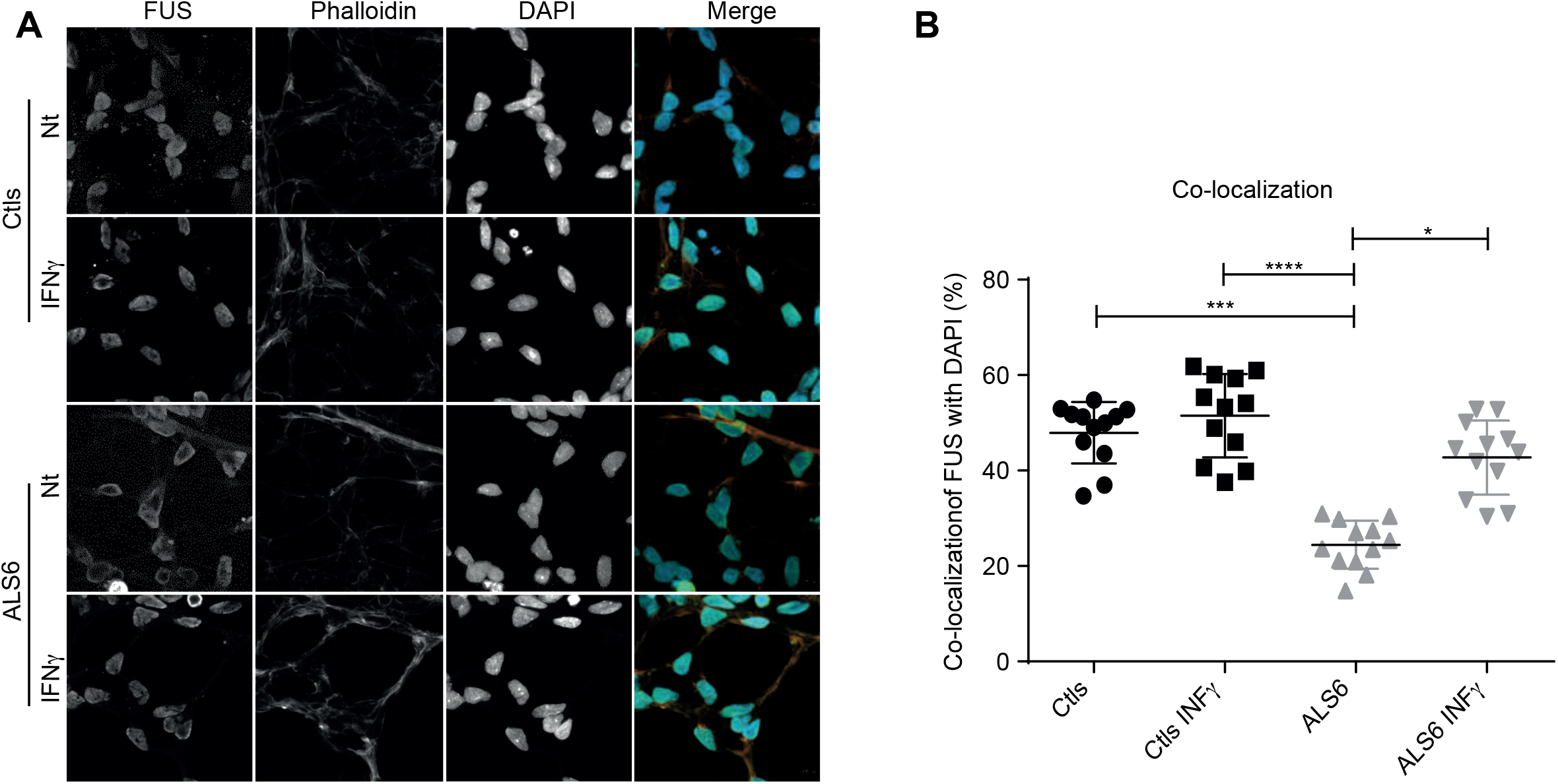
FUS localization in ALS6 cells is also recovered by IFNγ treatment. **A**. Representative images of immunofluorescence stainings for FUS protein in control-and IFNγ- treated MNs. Scale bar represents 10 μm. **B**. Quantification of FUS localization from immunofluorescence stainings for FUS protein in control- and IFNγ-treated MNs. * = p<0.05; ** = p<0.01; and *** = p<0.001, two-way ANOVA with Tukey multiple comparison test; n = 8 per group.

**Extended Table 1:** List of the proteins found to interact with only controls, both controls and ALS6 or only ALS6 cells in the shotgun proteomics experiments.

| Samples | Number of proteins | Protein Gene name |  |  |  |  |
| --- | --- | --- | --- | --- | --- | --- |
| Controls and ALS6 | 25 | YBX1 | IGF2BP3 | ACTG1 | HNRNPA1 | HNRNPC |
|  |  | RPS2 | SYNCRIP | HNRNPU | PABPC1 | RPL4 |
|  |  | RPS18 | HIST1H4A | ELAVL1 | HIST1H2BM | HIST1H2AJ |
|  |  | HNRNPH1 | RPS3 | RPS16 | UPF1 | DHX9 |
|  |  | FUS | VIM | RPS9 | IGF2BP1 | RPL31 |
| ALS6 | 58 | H2AFY2 | HNRNPA0 | NES | LMNB1 | RPL7A |
|  |  | RPS17 | IGF2BP2 | EEF1A1 | RPS4X | RALY |
|  |  | RPS3A | RPL6 | PTBP1 | PRPF8 | MSI2 |
|  |  | FXR1 | GNB2L1 | EIF3A | KHDRBS1 | HNRNPA3 |
|  |  | DDX5 | RPL3 | RPL13 | HNRNPM | ILF2 |
|  |  | RPSA | ILF3 | SAFB | RBMX | SFPQ |
|  |  | HNRNPR | RPS14 | RPS8 | HIST1H1B | DHX15 |
|  |  | H3F3A | RPL27 | HNRNPK | HIST1H1C | LMNB2 |
|  |  | G3BP1 | NONO | RPL23A | MATR3 | G3BP2 |
|  |  | HNRNPA2B1 | RPS7 | H2AFY | XRCC6 | RPS10 |
|  |  | HNRNPD | HIST2H2BE | SF3B3 | NPM1 | HSPA8 |
|  |  | RPLP0 | CAPRIN1 | RPS19 |  |  |
| Controls | 5 | RPS25 |  |  |  |  |
|  |  | RPL17 |  |  |  |  |
|  |  | RPS6 |  |  |  |  |
|  |  | RPL18 |  |  |  |  |
|  |  | RPS20 |  |  |  |  |

